# IFN signaling and neutrophil degranulation transcriptional signatures are induced during SARS-CoV-2 infection

**DOI:** 10.1101/2020.08.06.239798

**Authors:** Bruce A. Rosa, Mushtaq Ahmed, Dhiraj K. Singh, José Alberto Choreño-Parra, Journey Cole, Luis Armando Jiménez-Álvarez, Tatiana Sofía Rodríguez-Reyna, Bindu Singh, Olga Gonzalez, Ricardo Carrion, Larry S. Schlesinger, John Martin, Joaquín Zúñiga, Makedonka Mitreva, Shabaana A. Khader, Deepak Kaushal

**Author notes:** Equal authorship (Bruce A. Rosa, Mushtaq Ahmed, Dhiraj K. Singh). **Corresponding authors:** Deepak Kaushal, Southwest National Primate Research Center, Texas Biomedical Research Institute, San Antonio, TX 78245,; Shabaana A. Khader, Department of Molecular Microbiology, Washington University in St. Louis, St. Louis, MO 63110,; and Makedonka Mitreva, Department of Medicine, Washington University in St. Louis, St. Louis, MO 63110,.

## Abstract

The novel virus SARS-CoV-2 has infected more than 14 million people worldwide resulting in the Coronavirus disease 2019 (COVID-19). Limited information on the underlying immune mechanisms that drive disease or protection during COVID-19 severely hamper development of therapeutics and vaccines. Thus, the establishment of relevant animal models that mimic the pathobiology of the disease is urgent. Rhesus macaques infected with SARS-CoV-2 exhibit disease pathobiology similar to human COVID-19, thus serving as a relevant animal model. In the current study, we have characterized the transcriptional signatures induced in the lungs of juvenile and old rhesus macaques following SARS-CoV-2 infection. We show that genes associated with Interferon (IFN) signaling, neutrophil degranulation and innate immune pathways are significantly induced in macaque infected lungs, while pathways associated with collagen formation are downregulated. In COVID-19, increasing age is a significant risk factor for poor prognosis and increased mortality. We demonstrate that Type I IFN and Notch signaling pathways are significantly upregulated in lungs of juvenile infected macaques when compared with old infected macaques. These results are corroborated with increased peripheral neutrophil counts and neutrophil lymphocyte ratio in older individuals with COVID-19 disease. In contrast, pathways involving VEGF are downregulated in lungs of old infected macaques. Using samples from humans with SARS-CoV-2 infection and COVID-19, we validate a subset of our findings. Finally, neutrophil degranulation, innate immune system and IFN gamma signaling pathways are upregulated in both tuberculosis and COVID-19, two pulmonary diseases where neutrophils are associated with increased severity. Together, our transcriptomic studies have delineated disease pathways to improve our understanding of the immunopathogenesis of COVID-19 to facilitate the design of new therapeutics for COVID-19.

## INTRODUCTION

COVID-19, caused by the novel severe acute respiratory syndrome coronavirus 2 (SARS-CoV-2), emerged as a pandemic disease during the end of 2019 and beginning of 2020. In the absence of a specific treatment or vaccine against SARS-CoV-2, infected individuals develop symptoms associated with a cytokine storm (*1*). This cytokine storm can initiate viral sepsis and inflammation-induced lung injury which lead to other complications including pneumonitis, acute respiratory distress syndrome (ARDS), respiratory failure, shock, organ failure and potentially death (*1, 2*).

By combining established principles of anti-viral immunity with analysis of immune responses in COVID-19 patients, a picture of the host defense response against SARS-CoV-2 is beginning to emerge (*3, 4*). Upon infection of the mucosal epithelium, SARS-CoV-2 is detected by intracellular pattern recognition receptors (PRRs) that bind viral RNA and DNA. PRR signaling triggers activation of transcription factors and induces Interferon (IFN) signaling, which in turn activates resident macrophages. Infected macrophages induce cytokine secretion that consequently triggers recruitment of myeloid cells, likely resulting in a feed-back loop that aggravates immunopathogenesis and promotes disease progression.

Analyses of transcriptomic response of host cells upon virus infection have potential to identify the host immune response dynamics and gene activated regulatory networks (*5, 6*). Recent studies have reported transcriptional changes in cells in the broncho-alveolar lavage (BAL) and peripheral blood mononuclear cells (PBMCs) of COVID-19 patients (*7*). Single cell RNA-seq has recently identified initial cellular targets of SARS-CoV-2 infection in model organisms (*8*) and patients (*9*) and characterized peripheral and local immune responses in severe COVID-19 (*10*), with severe disease being associated with a cytokine storm and increased neutrophil accumulation. However, most of these studies have mostly been performed in peripheral blood samples from a limited number of moderate or severe COVID-19 patients within limited age ranges (*10*). To overcome the limitations associated with obtaining samples from human subjects and to get more in-depth understanding of the transcriptional changes during COVID-19, we have developed a SARS-CoV-2 macaque model, where both juvenile and old macaques were infected and exhibited clinical symptoms that reflect human COVID-19 disease that is self-limited. In the current study, we have characterized the transcriptional signatures induced in the lungs of juvenile and old rhesus macaques following SARS-CoV-2 infection. Our results show that genes associated with Interferon (IFN) signaling, neutrophil degranulation and innate immune pathways are significantly induced in the lungs in response to SARS-CoV-2 infection. Interestingly, this is associated with a downregulation of genes associated with collagen formation and regulation of collagen pathways. In COVID-19, increasing age is a significant risk factor for poor prognosis of infection(*11*). We demonstrate that specific immune pathways, namely Type I IFN and Notch signaling, are significantly upregulated in juvenile macaques when compared with old macaques infected with SARS-CoV-2. These results are corroborated with increased peripheral neutrophil counts and neutrophil lymphocyte ratio in older individuals with COVID-19 disease. In contrast, the VEGF pathway is downregulated in old infected macaques. Incidently, levels of VEGF protein are increased in plasma of older COVID-19 patients, emphasizing the importance of studying both local and peripheral responses. Finally, we report that neutrophil degranulation, innate immune system and IFN gamma (IFN-γ) signaling pathways are upregulated in both tuberculosis (TB) and COVID-19, two pulmonary infectious diseases where neutrophils accumulation is associated with increased severity. Together, our study has delineated disease pathways that can serve as a valuable tool in understanding the immunopathogenesis of SARS-CoV-2 infection and progressive COVID-19, and facilitate the design of therapeutics for COVID-19.

## MATERIALS AND METHODS

### Macaques

All of the infected animals were housed in Animal Biosafety Level 3 (ABSL3) at the Southwest National Primate Research Center, Texas Biomedical Research Institute, where they were treated per the standards recommended by AAALAC International and the NIH Guide for the Care and Use of Laboratory Animals. Sham controls were housed in ABSL2. The animal studies in each of the species were approved by the Animal Care and Use Committee of the Texas Biomedical Research Institute and as an omnibus Biosafety Committee protocol.

### Animal studies, and tissue harvest for RNA sample preparation

Rhesus macaques (*Macaca mulatta*) animals enrolled in this study have been described in detail(*12*) (in review), and the infection of these animals with 1.05×10^6^ pfu SARS-CoV-2 isolate USA-WA1/2020 (BEI Resources, NR-52281, Manassas, VA) has also been described earlier(*12*) (in review). Control (SARS-CoV-2 uninfected) samples were obtained from opportunistic necropsies conducted on rhesus macaques from the same colony in the past few months. Infected animals were euthanized for tissue collection at necropsy, including lung. specimens Lung tissue from three juvenile (3 yrs old) and five old (average 17 yrs old) rhesus macaques (**Table S1**) were homogenized, snap-frozen in RLT buffer, and DNAse-treated total RNA was extracted using the Qiagen RNeasy Mini kit (Qiagen) for RNA-seq analysis as described earlier(*13*).

### Viral RNA determination

SARS-CoV-2 RNA isolation and measurement of viral RNA in lung homogenates using RTqPCR has been described(*12*) (in review).

### RNA-sequencing and analysis

cDNA libraries were prepared from RNA samples using the Clontech SMARTer universal low input RNA kit to maximize yield, and samples were sequenced on Illumina NovaSeq S4 XP (paired 150bp reads). After adapter trimming using Trimmomatic v0.39(*14*), sequenced RNA-seq reads were aligned to the *Macaca mulatta* genome (version 10, Ensembl release 100(*15*)) using the STAR aligner v2.7.3a(*16*) (2-pass mode, basic). All raw RNA-Seq fastq files were uploaded to the NCBI Sequence Read Archive (SRA(*17*)), and complete sample metadata and accession information are provided in **Table S1**. Read fragments (read pairs or single reads) were quantified per gene per sample using featureCounts v1.5.1(*18*). Significantly differentially expressed genes between naïve, controller and progressor sample sets were identified using DESeq2 v1.4.5(*19*) with default settings, and a minimum P value significance threshold of 0.01 (after False Discovery Rate [FDR(*20*)] correction for the number of tests). Principal components analysis also was calculated using DESeq2 output (default settings, using the top 500 most variable genes). FPKM (fragments per kilobase of gene length per million reads mapped) normalization was performed using DESeq2-normalized read counts. Pathway enrichment analysis among differentially expressed gene sets of interest was performed for (a) Reactome(*21*) pathways, using the human orthologs as input into the WebGestalt(*22*) web server (p ≤ 0.05 after FDR correction, minimum 3 genes per term) and (b) KEGG(*23*) pathways and Gene Ontology(*24*) terms, using the g:profiler web server(*25*) which has a database of these annotations matched to macaque ENSEMBL gene IDs (p ≤ 0.05 after FDR correction, minimum 3 genes per term). Mapped fragment counts, relative gene expression levels, gene annotations, and differential expression data for every macaque gene are available in **Table S2**, along with orthology matches to human genes retrieved from ENSEMBL(*15*) and identifications of differentially expressed (DE) genes belonging to enriched pathways of interest, for genes of interest in **Table S3**, and significant functional enrichment for Reactome, KEGG and Gene Ontology pathways, among differentially gene sets of interest in **Table S4**. Additionally, genes significantly differentially regulated during progression of tuberculosis (in both the macaque gene and the corresponding mouse ortholog) were identified from a previous transcriptomic study of tuberculosis-infected lung tissue(*13*), and the upregulated and downregulated gene sets were intersected with the COVID-19 results from the current study.

### Human sample collection

Plasma samples were collected from COVID-19 patients that attended the emergency room of the Instituto Nacional de Ciencias Médicas y Nutrición Salvador Zubirán (INCMNSZ), and the Instituto Nacional de Enfermedades Respiratorias Ismael Cosío Villegas (INER) in Mexico City, from March to June of 2020. Detection of SARS-CoV-2 was performed by real-time polymerase chain reaction (RT-PCR) in swab samples, bronchial aspirates (BA), or bronchoalveolar lavage (BAL). For this purpose, viral RNA was extracted from clinical samples with the MagNA Pure 96 system (Roche, Penzberg, Germany). The RT-PCR reactions were performed in a total volume of 25 μL, containing 5μL of RNA, 12.5μL of 2 × reaction buffer provided with the Superscript III one-step RT-PCR system with Platinum Taq Polymerase (Invitrogen, Darmstadt, Germany; containing 0.4 mM of each deoxyribose triphosphates (dNTP) and 3.2 mM magnesium sulfate), 1μL of reverse transcriptase/ Taq mixture from the kit, 0.4 μL of a 50 mM magnesium sulfate solution (Invitrogen), and 1μg of nonacetylated bovine serum albumin (Roche). All oligonucleotides were synthesized and provided by Tib-Molbiol (Berlin, Germany). Thermal cycling was performed at 55 °C for 10 min for reverse transcription, followed by 95 °C for 3 min and then 45 cycles of 95°C for 15 s, 58°C for 30s. Primer and probe sequences are as follows: RdRP gene [RdRp-SARSr-F:GTGARATGGTCATGTGTGGCGG,RdRp-SARSr-P2:

FAMCAGGTGGAACCTCATCAGGAGATGCBBQ,RdRP_SARSrP1:FAMCCAGGTGGWACRTC ATCMGGTGATGCBBQ,RdRp_SARSrR:CARATGTTAAASACACTATTAGCATA], E gene [E_Sarbeco_F:ACAGGTACGTTAATAGTTAATAGCGT,E_Sarbeco_P1:FAMACACTAGCCATC CTTACTGCGCTTCGBBQ,E_Sarbeco_R:ATATTGCAGCAGTACGCACACA], N gene [N_Sarbeco_F:CACATTGGCACCCGCAATC,N_Sarbeco_P1:FAMACTTCCTCAAGGAACAACA TTGCCABBQ, N_Sarbeco_R:GAGGAACGAGAAGAGGCTTG]. Clinical and demographic data were retrieved from the medical records of all participants. These data included age, gender, anthropometrics, comorbidities, symptoms, triage vital signs, and initial laboratory test results. Initial laboratory tests were defined as the first test results available (typically within 24 h of admission) and included white blood cell counts (WBC), neutrophil and lymphocyte counts (**Table S5**).

### Cytokine levels in human plasma samples

Peripheral blood samples were obtained from all participants at hospital admission. Plasma levels of interferon-gamma (IFN-γ) and vascular endothelial growth factor (VEGF), were determined by Luminex assays using the Luminex platform Bio-Plex Multiplex 200 (Bio-Rad Laboratories, Inc., Hercules, CA, USA). Plasma samples from four healthy volunteer donors were used as controls.

## RESULTS

### Genes up-regulated in COVID-19 infected macaques represent pathways characteristic of neutrophil degranulation and IFN signaling

We recently assessed the ability of SARS-CoV-2 to infect rhesus macaques during a longitudinal two week infection study. This study included the effect of age on the progression of infection to COVID-19. Indian-origin, SPF-rhesus macaques (*Macaca mulatta*) were infected by multiple routes (ocular, intratracheal and intranasal) with sixth-passage virus at a target dose of 1.05×10^6^ PFU/per animal and studied for two weeks. The macaques were grouped as naïve (uninfected), and infected (juvenile or old) macaques. All infected animals developed clinical signs of viral infection(*12*) (in review). Both juvenile and old macaques exhibited comparable clinical disease, and equivalent longitudinal viral loads in the BAL, nasopharyngeal and buccopharyngeal swabs, as well as lungs at endpoint. This was followed by comparable viral clearance. In order to fully understand the immune pathways regulated upon SARS-CoV-2 infection, RNA was extracted and RNA sequencing was carried out from a lung biopsy from juvenile macaques (n = 3, 1 male and 2 females) and old macaques infected with infected with SARS-CoV-2 (n = 5, 1 male and 4 females) and naive uninfected macaques (n = 4, 2 males and 2 females). An average of 68.6 million reads were generated, with an average of 20.3 million fragments (read pairs or orphaned reads) mapping to macaque coding sequences, following analytical processing and mapping (**Table S1**). Principal components analysis (PCA) based on whole-transcriptome gene expression levels(*19*) showed that despite within-group variability for the COVID-19 infected samples, the naive samples grouped separately, suggesting substantial overall transcriptomic differences resulting from the infection (**Figure 1A**). Differential gene expression analysis (DESeq2(*19*)) with the juvenile and old COVID-19 samples grouped together identified 1,026 genes significantly (P ≤ 0.01) up-regulated in response to infection, while 1,109 genes were significantly downregulated (**Figure 1B**). Expression, annotation and differential expression data for all genes is available in **Table S2**. Complete lists of differentially expressed genes for each comparison of interest (described below) ranked by P value, with Z-scores for expression visualization are available in **Table S3**, and significant pathway enrichment (Reactome(*21*), KEGG(*23*) and Gene Ontology(*24*)) for all comparisons is shown in **Table S4**.

**Figure 1:**
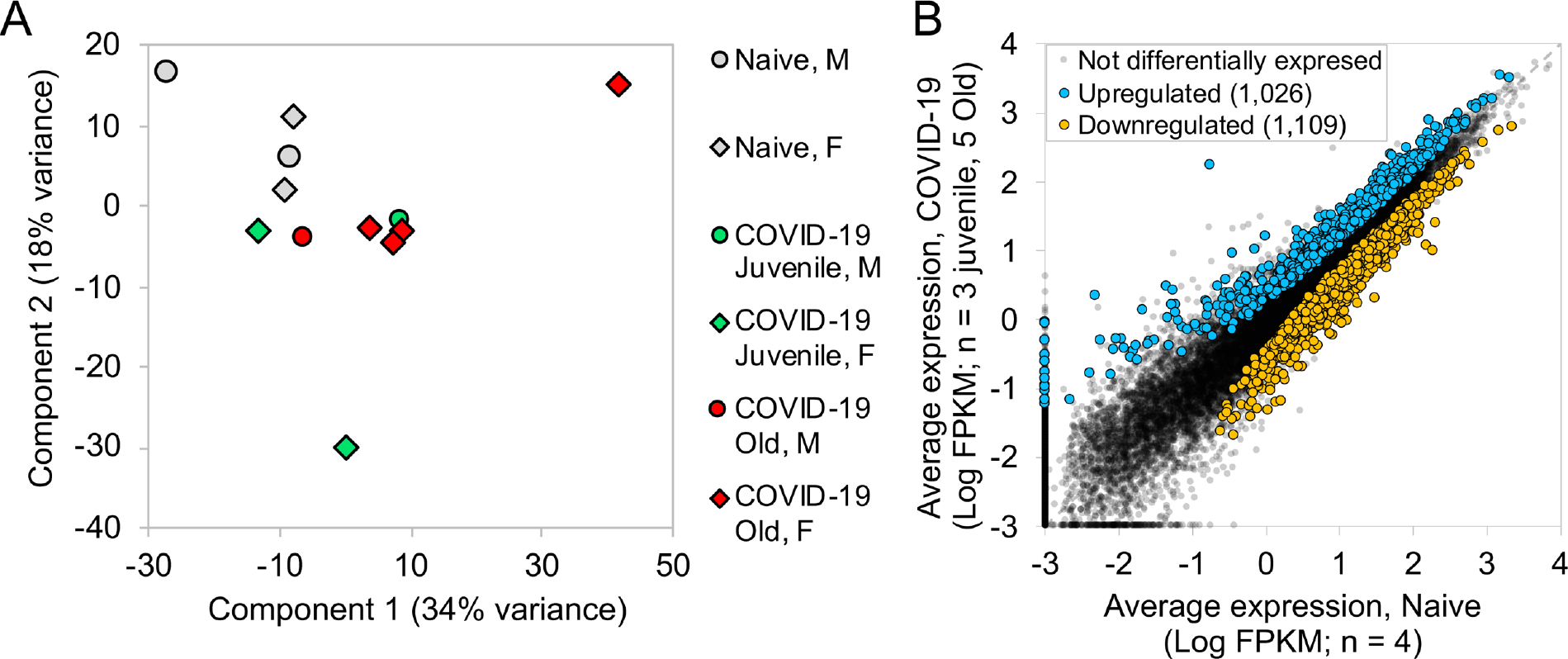
Genes upregulated in COVID-19-infected macaques represent pathways characteristic of neutrophil degranulation and IFN signaling. Differential gene expression between naive and COVID-19 samples. (**A**) PCA plot showing the clustering of samples based on overall transcriptomic profiles. (**B**) Gene expression plot showing the relative normalized gene expression levels (FPKM) for each gene, with genes significantly differentially regulated by COVID-19 indicated.

Evaluation of the top 30 most significantly up-regulated genes in the lungs of SARS-CoV-2-infected macaques revealed significantly higher expression of CTSG (Cathepsin G), ATP6AP2(ATPase H+ transporting accessory protein 2), IFNγR1 (Interferon Gamma Receptor), CD36 and CD58, in comparison to expression in uninfected macaque lungs (**Figure 2A**). Cathepsin G is a serine protease prominently found in neutrophilic granules. IFNγR1 associates with IFNγR2 to form a receptor for the cytokine interferon gamma (IFNγ)(*26–29*), and required for activation of antiviral responses, such as IRF3 (IFN regulatory factor-3), nuclear factor KB (NF-KB) and JAK (Janus kinase)/STAT (signal transducer and activator of transcription) signaling pathways (*30*). Reactome pathway analysis on up- and down-regulated genes in the lungs of SARS-CoV-2 infected rhesus macaques showed that genes significantly up-regulated by infection, included pathway enrichment for genes involved in “Neutrophil degranulation”, “Innate Immune system”, “Immune system” and “IFN signaling” (**Table 1**; **Table S4A**). The up-regulation of CD36 during COVID-19 in lungs is in conformity with these enriched pathways, since CD36, a scavenger receptor expressed in multiple cell types, mediates lipid uptake, immunological recognition, inflammation, molecular adhesion, and apoptosis (*31*), and is a Matrix Metalloproteinase-9 substrate that induces neutrophil apoptosis. CD58 molecule (lymphocyte function-associated antigen-3) is expressed on human hematopoietic and non-hematopoietic cells, including dendritic cells, macrophages and endothelial cells (*32–35*), and interacts with its receptor CD2 molecule (*36, 37*) on CD8^+^ cytotoxic T lymphocytes and NK cells to mediate cytotoxic reactions (*38–40*). The complete ranked list of the 1,026 genes upregulated during COVID-19 is shown in **Table S3A**.

**Figure 2:**
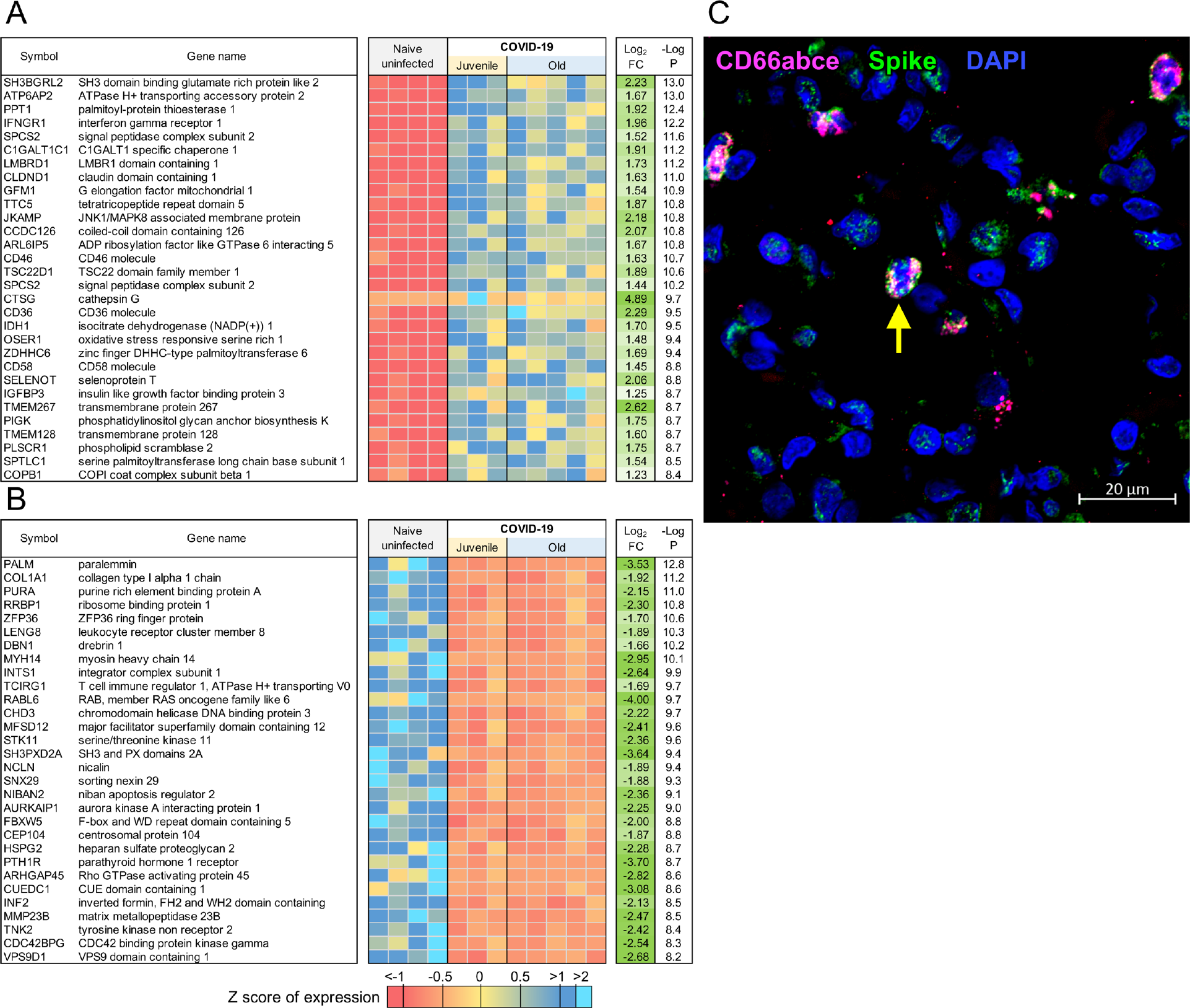
Genes downregulated in COVID-19-infected macaques represent pathways characteristic of collagen degradation and TFG-β signaling. The top 30 most significantly (**A**) upregulated genes and (**B**) downregulated genes in COVID-19 infected macaque lungs. Expression values are visualized by Z scores of normalized expression data (FPKM) per sample, and Log2 Fold Change and −Log P values are from the DESeq2 output. Genes are sorted by P value. (**C**) Multilabel confocal immunofluorescence microscopy of FFPE lung sections from SARS CoV-2 infected rhesus macaques with SARS CoV-2 Spike specific antibody (green), neutrophil marker CD66abce (red) and DAPI (blue) at 10X magnification.

**Table 1:**
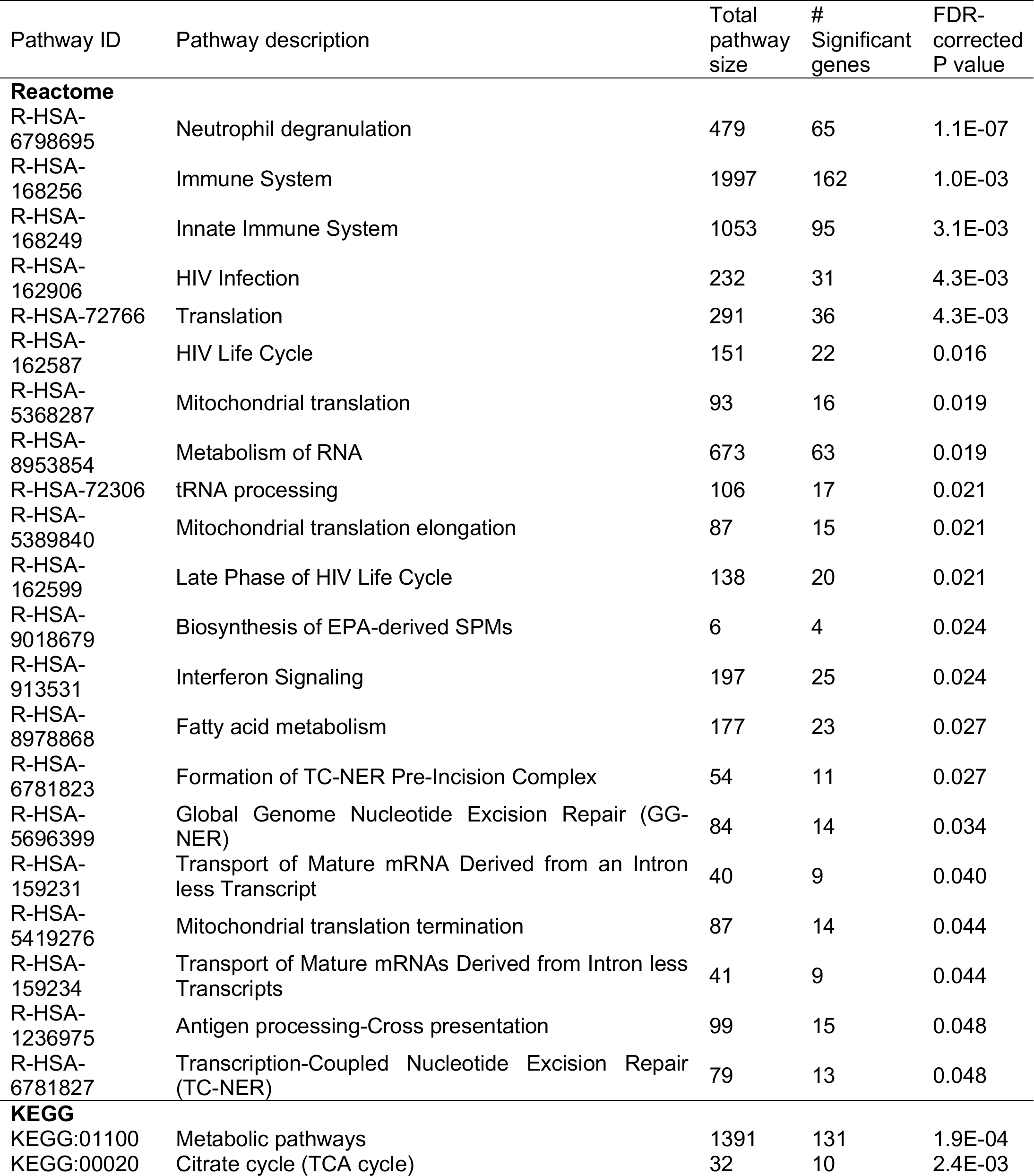
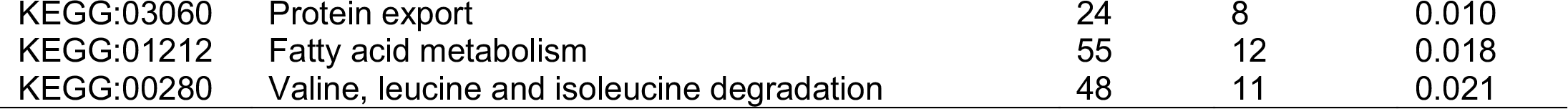
Significant Reactome and KEGG pathway enrichment among the 1,026 genes significantly upregulated by COVID-19.

| Pathway ID | Pathway description | Total pathway size | # Significant genes | FDR-corrected P value |
| --- | --- | --- | --- | --- |
| **Reactome** |  |  |  |  |
| R-HSA-6798695 | Neutrophil degranulation | 479 | 65 | 1.1E-07 |
| R-HSA-168256 | Immune System | 1997 | 162 | 1.0E-03 |
| R-HSA-168249 | Innate Immune System | 1053 | 95 | 3.1E-03 |
| R-HSA-162906 | HIV Infection | 232 | 31 | 4.3E-03 |
| R-HSA-72766 | Translation | 291 | 36 | 4.3E-03 |
| R-HSA-162587 | HIV Life Cycle | 151 | 22 | 0.016 |
| R-HSA-5368287 | Mitochondrial translation | 93 | 16 | 0.019 |
| R-HSA-8953854 | Metabolism of RNA | 673 | 63 | 0.019 |
| R-HSA-72306 | tRNA processing | 106 | 17 | 0.021 |
| R-HSA-5389840 | Mitochondrial translation elongation | 87 | 15 | 0.021 |
| R-HSA-162599 | Late Phase of HIV Life Cycle | 138 | 20 | 0.021 |
| R-HSA-9018679 | Biosynthesis of EPA-derived SPMs | 6 | 4 | 0.024 |
| R-HSA-913531 | Interferon Signaling | 197 | 25 | 0.024 |
| R-HSA-8978868 | Fatty acid metabolism | 177 | 23 | 0.027 |
| R-HSA-6781823 | Formation of TC-NER Pre-Incision Complex | 54 | 11 | 0.027 |
| R-HSA-5696399 | Global Genome Nucleotide Excision Repair (GG-NER) | 84 | 14 | 0.034 |
| R-HSA-159231 | Transport of Mature mRNA Derived from an Intron less Transcript | 40 | 9 | 0.040 |
| R-HSA-5419276 | Mitochondrial translation termination | 87 | 14 | 0.044 |
| R-HSA-159234 | Transport of Mature mRNAs Derived from Intron less Transcripts | 41 | 9 | 0.044 |
| R-HSA-1236975 | Antigen processing-Cross presentation | 99 | 15 | 0.048 |
| R-HSA-6781827 | Transcription-Coupled Nucleotide Excision Repair (TC-NER) | 79 | 13 | 0.048 |
| **KEGG** |  |  |  |  |
| KEGG:01100 | Metabolic pathways | 1391 | 131 | 1.9E-04 |
| KEGG:00020 | Citrate cycle (TCA cycle) | 32 | 10 | 2.4E-03 |
| KEGG:03060 | Protein export | 24 | 8 | 0.010 |
| KEGG:01212 | Fatty acid metabolism | 55 | 12 | 0.018 |
| KEGG:00280 | Valine, leucine and isoleucine degradation | 48 | 11 | 0.021 |

ATP6AP2 was the most significantly up-regulated of the 65 genes upregulated within the enriched “neutrophil degranulation” (R-HSA-6798695) pathway (**Table S3B**), and it interacts with renin or prorenin to cause activation of intracellular signaling pathways, resulting in secretion of inflammatory and fibrotic factors(*41*). CEACAM8 (Carcinoembryonic Antigen-Related Cell Adhesion Molecule 8) is the gene that encodes for CD66b, a well characterized marker of degranulation(*42*). Indeed, CD66b^+^ neutrophils accumulate in the lungs of macaques infected with SARS-CoV-2 (**Figure 2C**). We have also previously demonstrated that neutrophils are heavily recruited early to the alveolar space following SARS-CoV-2 infection of macaques(*12*) (in review). Additional genes strongly up-regulated during COVID-19 in the neutrophil degranulation pathway are IDH-1(Isocitrate Dehydrogenase (NADP(+)) 1) which regulates neutrophil chemotaxis, and FPR2 (Formyl Peptide Receptor 2), a G-coupled surface receptor which has a deleterious role to play in viral infection including influenza (*43*). LTA4H (Leukotriene A4 hydrolase) is an enzyme that generates a neutrophil chemoattractant, leukotriene B4, a marker for ARDS(*44*). Expression of 162 genes belonging to the “immune system” (R-HSA-168256) pathway was upregulated in SARS-CoV-2 infected macaques (**Table S3C**). These included LAMP-2(Lysosomal Associated Membrane Protein 2), and ATG7 (Autophagy Related 7), key genes involved in autophagy. LAMP-2 is known to influence phagosomal maturation in neutrophil (*45*). The IFN response constitutes the major first line of defense against viruses. Consistent with this, we found up-regulation of genes associated with the IFN signaling pathways, specifically Interferon Induced Protein with Tetratricopeptide Repeats 1 (IFIT3), IFN alpha receptor 1 (IFNAR1), IFN gamma receptor 1 (IFNGR1) and OAS 1 protein (2’-5’-Oligoadenylate Synthetase 1). Together, these results suggest that upregulation of neutrophil degranulation, Type I IFN signaling, and innate immune system is a characteristic feature of host responses to SARS-CoV-2 infection.

### Genes down-regulated following SARS-CoV-2 infection in macaques represent pathways characteristic of collagen degradation and TFG-β signaling

It is thought that up to 40% of patients with COVID-19 develop ARDS, and 20% of ARDS cases are severe (*46*). A well-documented sequela of ARDS is the development of fibrotic disease (*47, 48*). We found that the 1,109 genes downegulated in SARS-CoV-2-infected macaques were significantly enriched for collagen degradation, regulation and formation (**Figure 2B**; **Table 2**; **Table S3D**; **Table S4B**). For example, among the “collagen degradation” (R-HSA-1442490) enriched pathway (**Table S3E**), COLA1 (collagen type I chain), other members of the collagen gene family (COL4A2 COL16A1 COL4A4 COL6A2 COL6A1 COL5A1 COL9A1 COL13A1 COL12A1 COL1A2) and Matrix metalloproteases such as MMP23B (Matrix Metallopeptidase 23B), MMP15 and MMP14 were all significantly dowregulated in COVID-19 diseased lungs when compared with expression in lungs of uninfected controls. Additionally, Reactome pathway enrichment prominently featured pathways down-regulated in COVID-19 disease in macaques comprised of “collagen degradation”, “collagen chain trimerization”, “degradation of extracellular matrix” and “collagen formation” (**Table 2**). Increased collagen degradation is essential for the prevention of fibrosis, a sequelae of COVID-19 and ARDS. Therefore, regulation of collagen degradation and extracellular matrix modeling suggest that this may be a feature of SARS-CoV-2 infection of rhesus macaques being a self-limiting model with early and robust anamnestic responses. TGFβ (Transforming Growth Factor Beta 1) is involved in normal tissue repair following lung injury, and in mediating fibrotic tissue remodeling by increasing the production and decreasing the degradation of connective tissue (*49*). Our results indicate a downregulation of genes associated with TGFβ signaling (**Table 2**), including the genes PARD3 (par-3 family cell polarity regulator) and PARD6A (par-6 family cell polarity regulator alpha), which are involved in regulating epithelial cell apico-basolateral polarization, SMURF (SMAD specific E3 ubiquitin protein ligase 1), a negative regulator of TGFβ pathway, and FURIN, which is a TGFβ converting enzyme (**Table S3F**). While the interaction of the genes within these pathways is complex, our results project a broad downregulation of mechanisms that contribute to lung repair and remodeling in animals with anamnestic control of SARS-CoV-2 infection.

**Table 2:** Significant Reactome and KEGG pathway enrichment among the 1,109 genes significantly downregulated by COVID-19.

| Pathway ID | Pathway description | Total pathway size | # Significant genes | FDR-corrected P value |
| --- | --- | --- | --- | --- |
| **Reactome** |  |  |  |  |
| R-HSA-5653656 | Vesicle-mediated transport | 667 | 74 | 2.5E-05 |
| R-HSA-199991 | Membrane Trafficking | 628 | 66 | 6.1E-04 |
| R-HSA-1442490 | Collagen degradation | 64 | 14 | 6.4E-03 |
| R-HSA-73887 | Death Receptor Signaling | 141 | 22 | 6.4E-03 |
| R-HSA-194315 | Signaling by Rho GTPases | 444 | 47 | 7.8E-03 |
| R-HSA-8948216 | Collagen chain trimerization | 44 | 11 | 7.8E-03 |
| R-HSA-2022090 | Assembly of collagen fibrils and other multimeric structures | 61 | 13 | 7.9E-03 |
| R-HSA-1474290 | Collagen formation | 90 | 16 | 9.5E-03 |
| R-HSA-170834 | Signaling by TGF-beta Receptor Complex | 73 | 14 | 0.011 |
| R-HSA-1650814 | Collagen biosynthesis and modifying enzymes | 67 | 13 | 0.013 |
| R-HSA-3247509 | Chromatin modifying enzymes | 275 | 32 | 0.013 |
| R-HSA-4839726 | Chromatin organization | 275 | 32 | 0.013 |
| R-HSA-194840 | Rho GTPase cycle | 138 | 20 | 0.014 |
| R-HSA-2214320 | Anchoring fibril formation | 15 | 6 | 0.014 |
| R-HSA-9006934 | Signaling by Receptor Tyrosine Kinases | 455 | 45 | 0.022 |
| R-HSA-9006936 | Signaling by TGF-beta family members | 102 | 16 | 0.022 |
| R-HSA-446353 | Cell-extracellular matrix interactions | 18 | 6 | 0.033 |
| R-HSA-2243919 | Crosslinking of collagen fibrils | 18 | 6 | 0.033 |
| R-HSA-1474228 | Degradation of the extracellular matrix | 140 | 19 | 0.033 |
| R-HSA-193704 | p75 NTR receptor-mediated signaling | 97 | 15 | 0.033 |
| R-HSA-193648 | NRAGE signals death through JNK | 59 | 11 | 0.037 |
| R-HSA-416482 | G alpha (12/13) signaling events | 79 | 13 | 0.038 |
| R-HSA-3000480 | Scavenging by Class A Receptors | 19 | 6 | 0.038 |
| R-HSA-5140745 | WNT5A-dependent internalization of FZD2, FZD5 and ROR2 | 13 | 5 | 0.040 |
| R-HSA-9007101 | Rab regulation of trafficking | 124 | 17 | 0.047 |
| **KEGG** |  |  |  |  |
| KEGG:04144 | Endocytosis | 231 | 37 | 3.0E-06 |
| KEGG:05165 | Human papillomavirus infection | 314 | 42 | 6.3E-05 |
| KEGG:04510 | Focal adhesion | 194 | 29 | 4.3E-04 |
| KEGG:04530 | Tight junction | 150 | 24 | 9.1E-04 |
| KEGG:05135 | Yersinia infection | 116 | 20 | 1.7E-03 |
| KEGG:05132 | Salmonella infection | 205 | 26 | 0.023 |
| KEGG:04390 | Hippo signaling pathway | 146 | 20 | 0.045 |

### Type I interferon signaling and Notch signaling pathways are upregulated in young macaques but not old macaques with COVID-19 disease

Age is a significant risk factor for increased morbidity and mortality in COVID-19 disease (*11*). In order to identify the differential immune responses associated with SARS-CoV-2 infection in old macaques, we carried out differential expression analysis between the groups; namely between juvenile (n=3) vs naive (n=4), and old (n=5) vs naive (n=4). In order for a gene to be considered to be differentially expressed only in the juvenile macaques, we required a stringent P value for significance ≤ 0.01 in the juvenile COVID-19 vs naive, and a P value for significance ≥ 0.1 in the old COVID-19 vs naive comparison. This approach identified 86 genes significantly up-regulated (**Figure 3A**; **Table S3G**) and 96 genes significantly down-regulated (**Figure 3B**; **Table S3H**) with COVID-19 disease only in juveniles. Note that no genes were significantly upregulated in juveniles and significantly downregulated in old, and vice-versa. Of these genes, the top 30 most significantly differential between juvenile and old are shown for up-regulated genes in **Figure 4A** and for down-regulated genes in **Figure 4B**. No pathways were found to be significantly enriched among the 96 genes significantly downregulated only in juveniles, but the Reactome and KEGG pathways significantly enriched among the 86 genes upregulated only in juveniles are shown in **Table 3**. Complete gene lists per pathway, and all significant pathways enrichment results including for Gene Ontology (GO) are available in **Table S4C**.

**Figure 3:**
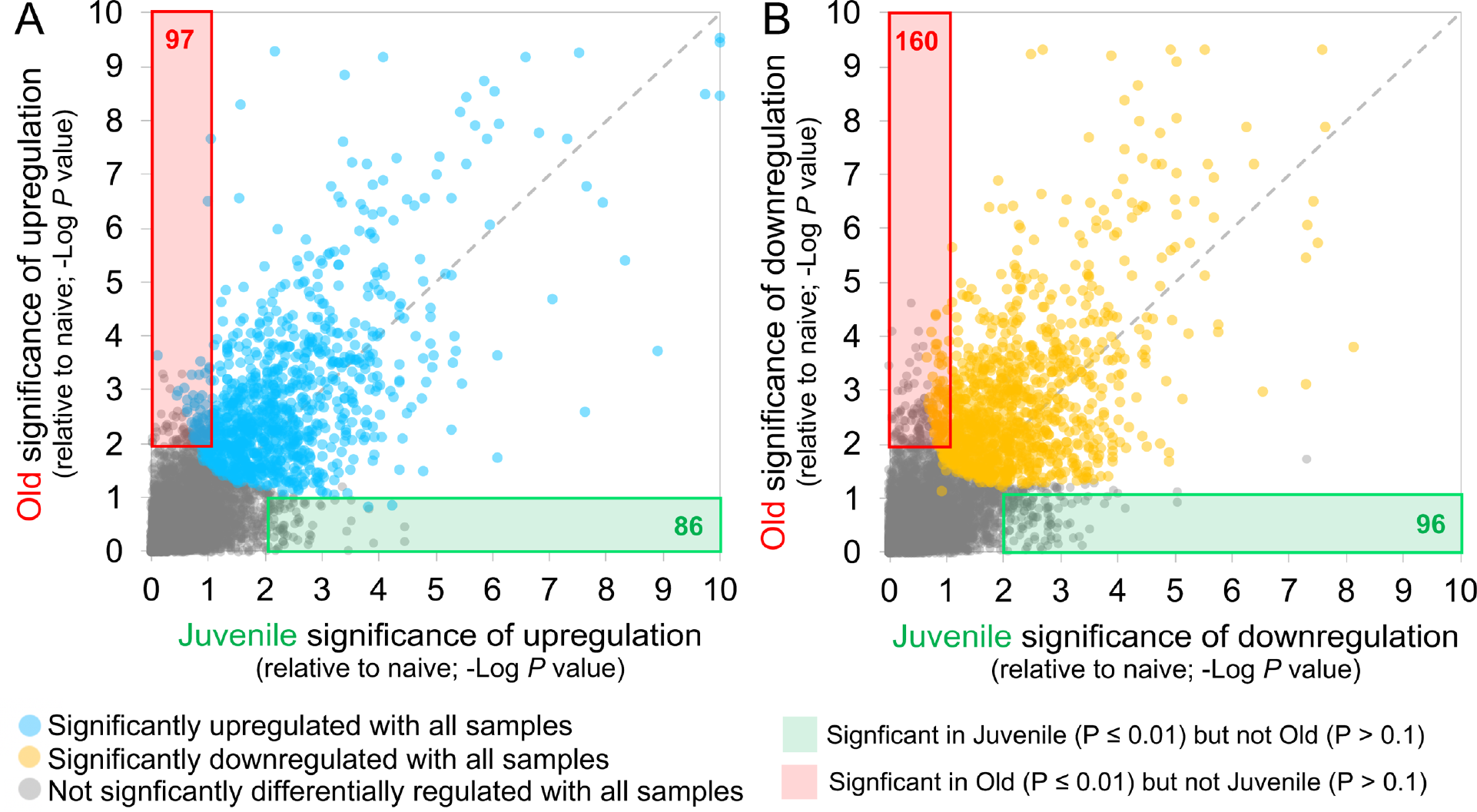
86 genes significantly upregulated and 96 genes significantly downregulated with COVID-19 only in juvenile macaques. Scatterplots visualizing the significance values of COVID-19 upregulated (**A**) and downregulated (**B**) genes, in juvenile and old macaques. Green shaded areas contain genes significant only in juveniles, and red shaded areas contain genes significant only in old macaques.

**Figure 4:**
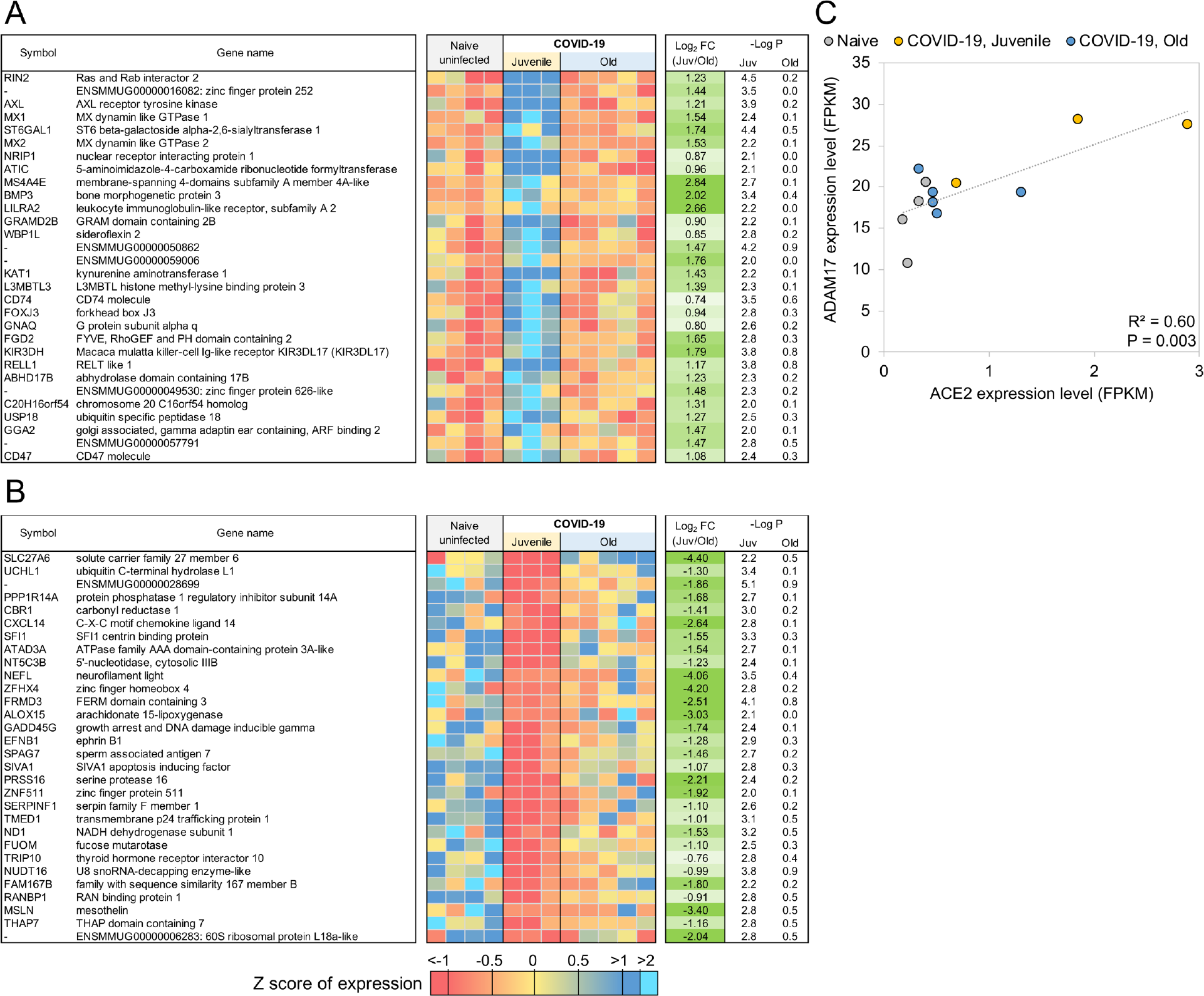
Genes related to Type I interferon signaling are upregulated in juvenile macaques compared to old macaques during COVID-19-infection. The top 30 most significantly (**A**) upregulated genes and (**B**) downregulated genes in COVID-19 infected juvenile macaque lungs but not in old macaques. Expression values are visualized by Z scores of normalized expression data (FPKM) per sample, and Log2 Fold Change and −Log P values are from the DESeq2 output. Genes are sorted by P value. (**C**) The relative gene expression of ACE2 and ADAM17 among naive, juvenile and old COVID-19 infected macaques.

**Table 3:**
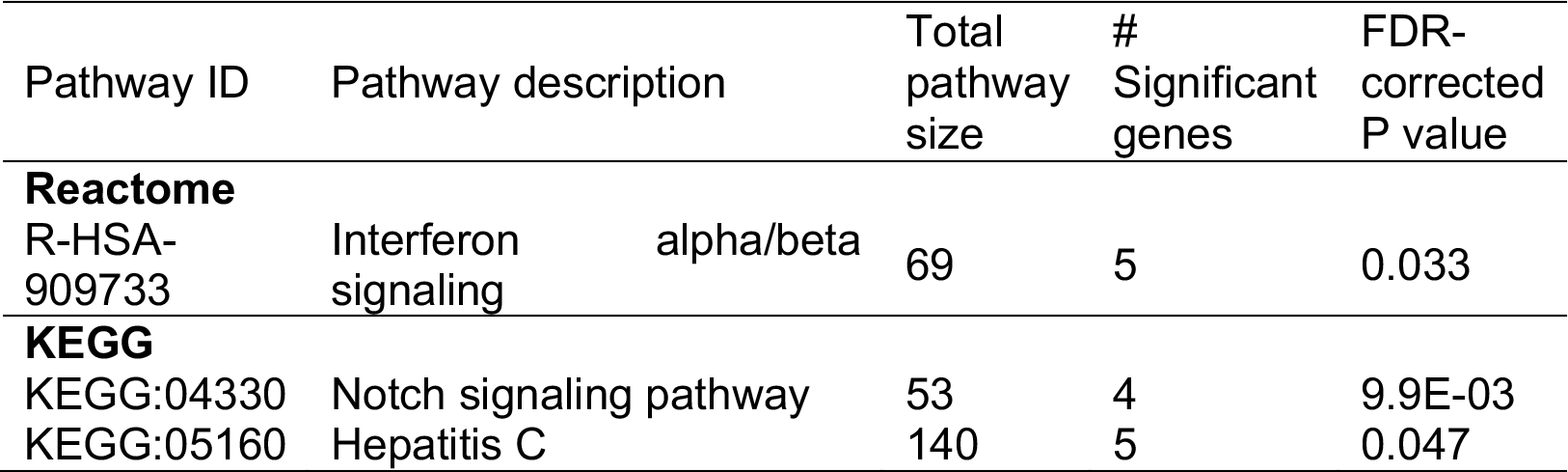
Significant Reactome and KEGG pathway enrichment among the 86 genes significantly upregulated by COVID-19 only in Juvenile macaques.

The genes with significantly upregulated expression in SARS-CoV-2 infected juvenile but not old macaques included MX1 (MX Dynamin Like GTPase 1), MX2 (MX Dynamin Like GTPase 2) and USP18 (Ubiquitin Specific Peptidase 18) (**Figure 5**). This is consistent with and highlights the role of the Reactome pathway “interferon alpha/beta signaling” being enriched in juvenile macaques during SARS-CoV-2 infection (**Table 3, Table S4C**). Other genes in this pathway which exhibited increased expression included IFIT1 and IFIT2. Additionally, by KEGG analysis, the Notch signaling pathway was observed to be significantly upregulated in juvenile infected macaques when compared with old infected macaques. ADAM17 (ADAM Metallopeptidase Domain 17), a key component of the Notch signaling pathways is known to be involved in shedding of the surface protein ACE2 (Angiotensin converting enzyme 2) (*50*). Therefore, it is interesting that a linear correlation in the expression of ACE2 and ADAM17 exists in infected macaques (**Figure 4C**). Note that we also see a significant upregulation of ACE2 across all samples (4.2-fold, P = 4.9×10^-3^), and a substantially larger upregulation among the juvenile samples (7.1-fold, P = 3.4×10^-4^). Additionally, the induction of DLL4, a Notch ligand, was increased in the infected juvenile macaques. Finally, the differential induction of DTX3L (Deltex E3 Ubiquitin Ligase 3L) in juvenile infected macaques compared to old infected macaques is important because Deltex stabilizes the receptor in the endocytic compartment allowing signal transduction to proceed in Notch signaling(*52*). Of the Hepatitis-induced pathway genes that are upregulated in juvenile COVID-19 diseased lungs, CXCL-10 (C-X-C Motif Chemokine Ligand 10) is a chemokine associated with severe disease in COVID-19 in humans (*53*), but can also be involved in recruitment of CXCR3 (C-X-C Motif Chemokine Receptor 3) expressing immune cells. 14-3-3 (otherwise called YWHAG) interacts with MDA5 (melanoma differentiation-associated protein 5), which belongs to the RIG-I-like receptor family and drive anti-viral immunity. Together, these results suggest that specific pathways including Type I IFN and Notch signaling are highly induced in juvenile macaques during SARS-CoV-2 infection, when compared to similarly infected old macaques.

**Figure 5:**
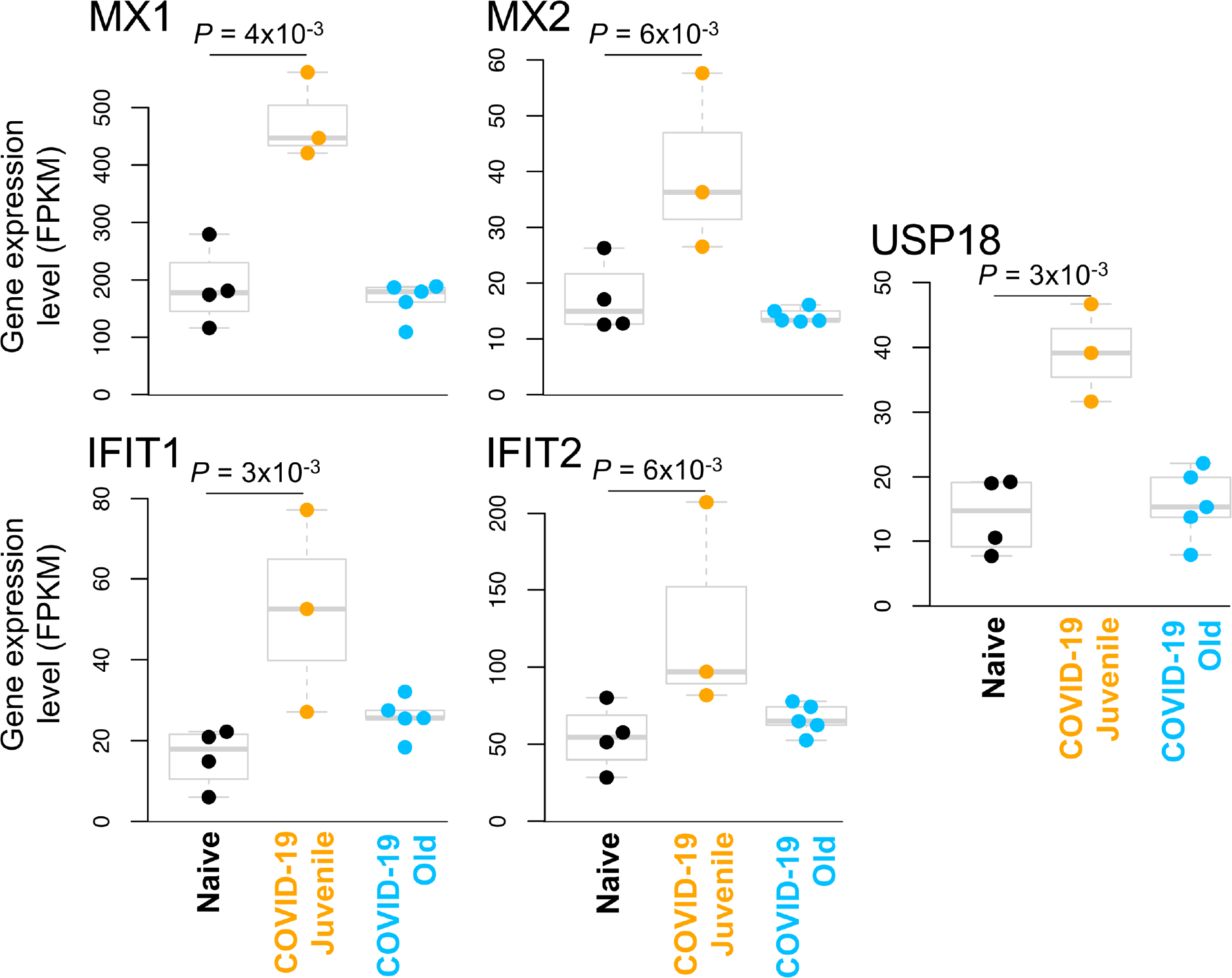
Interferon alpha signaling genes are significantly upregulated in juvenile COVID-19-infected macaques but not old COVID-19-infected macaques. The relative expression levels (FPKM) for the five “interferon alpha signaling” (HSA-909733) genes belonging to this gene set are shown. P values represent FDR-corrected significance values from DESeq2.

### Genes related to VEGF signaling are downregulated in old macaques but not juvenile macaques during COVID-19-disease

Using the same approach as for the juvenile macaque-specific differentially regulated genes, we identified 97 genes significantly up-regulated (**Figure 3A**; **Table S3I**) and 160 genes significantly down-regulated (**Figure 3B**; **Table S3J**) with COVID-19 disease only in infected old macaques, and not infected juveniles. Pathway enrichment analysis only identified significant functional enrichment among the down-regulated gene set (**Table 4**; **Table S4D**). Our results show that in the lungs of old macaques, the only Reactome pathways enriched among genes downregulated during COVID-19 included genes involved in the “VEGF-VEGFR2 Pathway” and “Signaling by VEGF” (**Figure 6, 7**). Vascular endothelial growth factor (VEGF) is a signaling protein that promotes angiogenesis, and is a key factor that promotes ARDS. Previous research showed that ACE2 antagonizes and down-regulates VEGFA(*54*), improving lung function following acute lung injury (*55*). Here, we observe both a significant increase in ACE2 in response to COVID-19 and a significant decrease in VEGF pathways in old macaques, which may be due to this antagonistic relationship. VEGFA, p21-activated kinase (PAK2), cytoplasmic tyrosine kinase (SRC), RhoA/ROCK signaling [ROCK1(Rho Associated Coiled-Coil Containing Protein Kinase 1) and WASF2(WASP Family Member 2) are all essential for multiple aspects of VEGF-mediated angiogenesis and are all significantly downregulated in old macaques with COVID-19 (**Figure 7**). Overall, despite juvenile and old macaques having a comparable clinical course with resolution, our data suggest that there are significant differenes in signaling pathways, especially those related to VEGF signaling that may ultimately result in differences is long term outcomes. Thus, our results suggest that down-regulation of VEGF pathways is associated with increasing age, in a macaque cohort of self-limiting disease model, and protect from serious lung injury during COVID-19 disease.

**Figure 6:**
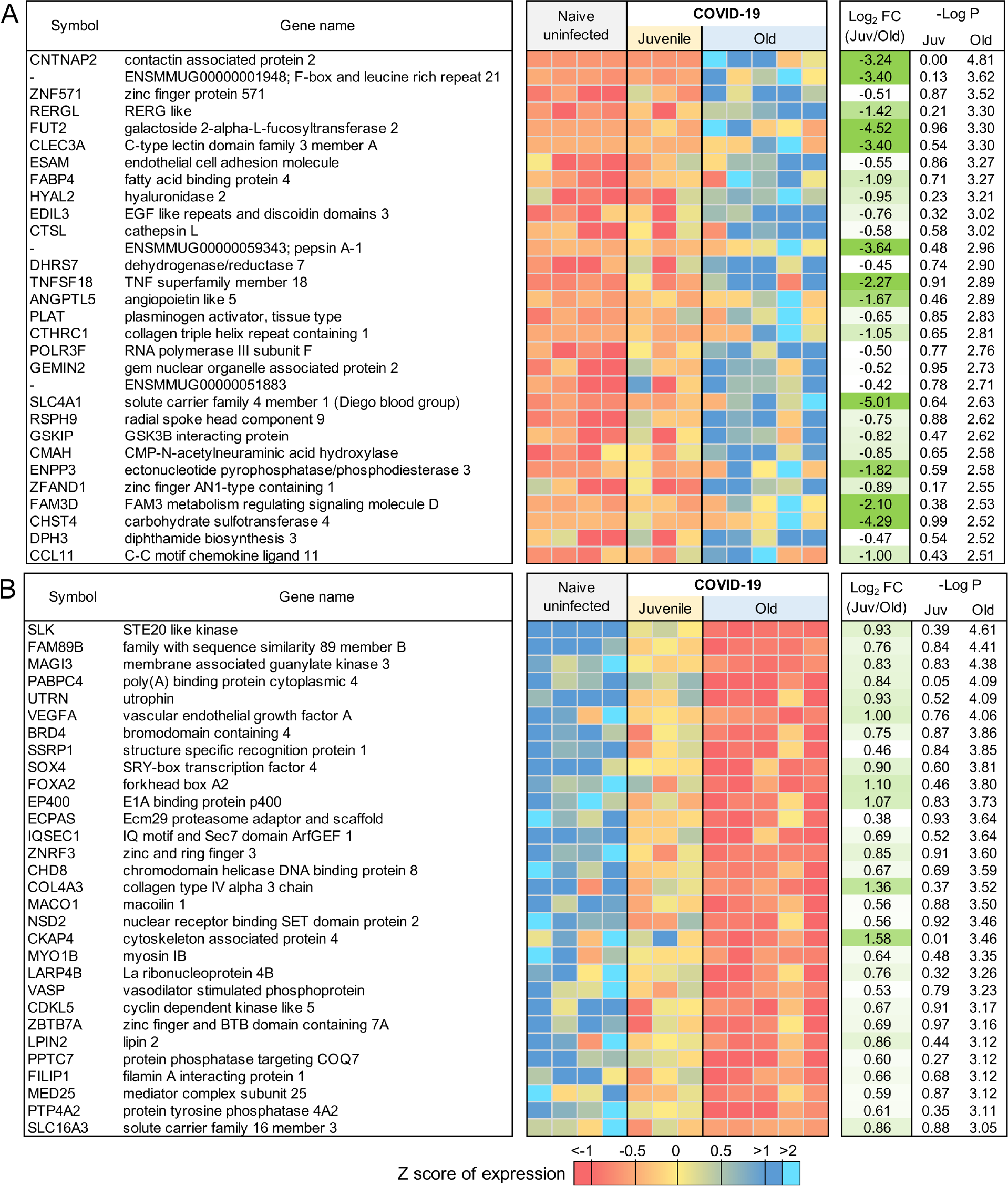
Genes related to VEGF signaling are downregulated in old macaques compared to juvenile macaques during COVID-19. The top 30 most significantly (**A**) upregulated genes and (**B**) downregulated genes in infected old macaque lungs but not in juvenile macaques. Expression values are visualized by Z scores of normalized expression data (FPKM) per sample, and Log2 Fold Change and −Log P values are from the DESeq2 output. Genes are sorted by P value.

**Figure 7:**
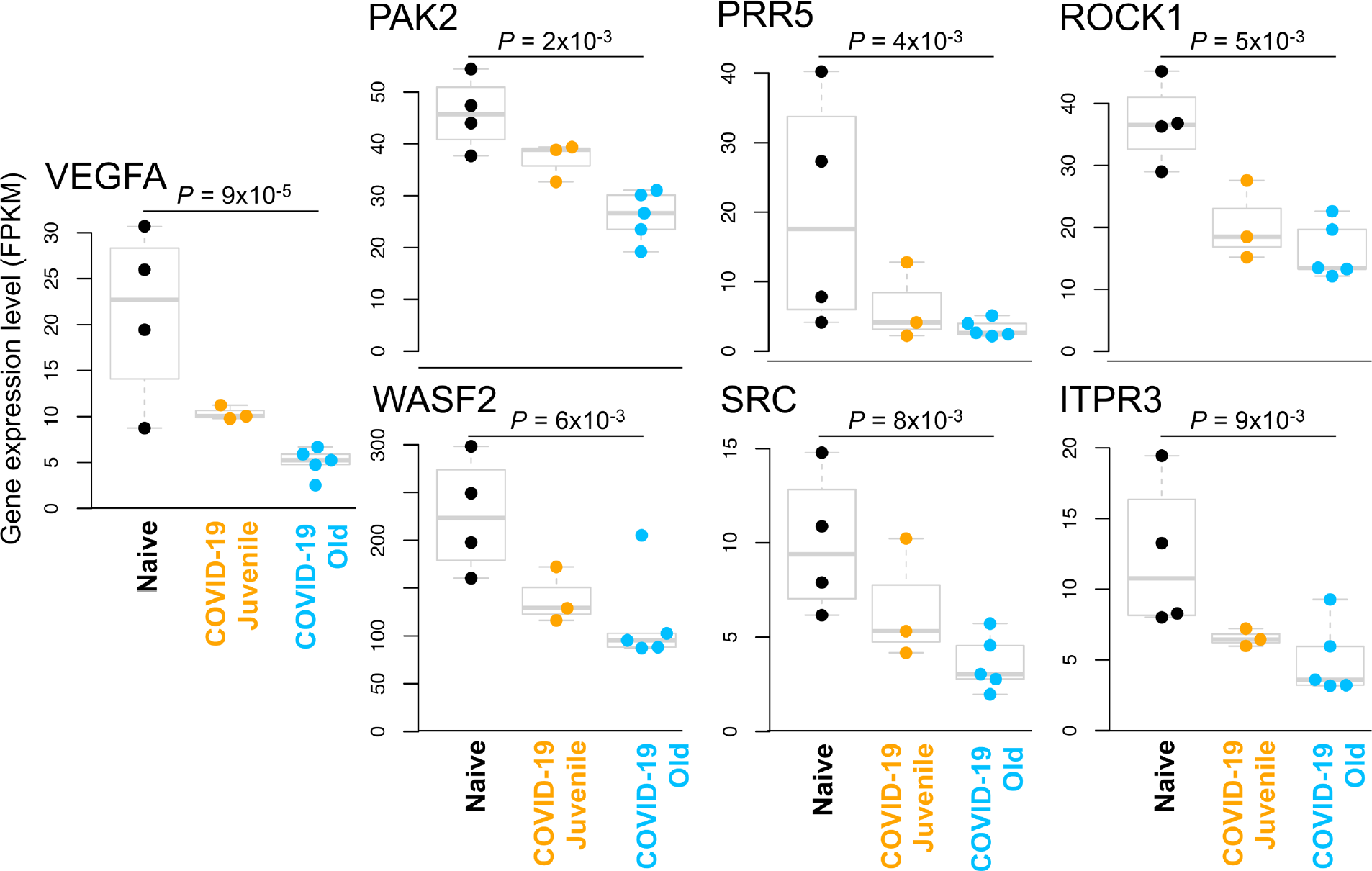
VEGF pathway genes are significantly downregulated in old COVID-19-infected macaques but not juvenile COVID-19-infected macaques. The relative expression levels (FPKM) for the seven “Signaling by VEGF” (R-HSA-194138) genes belonging to this gene set are shown. P values represent FDR-corrected significance values from DESeq2.

**Table 4:** Significant Reactome and KEGG pathway enrichment among the 160 genes significantly downregulated by COVID-19 only in Old macaques.

| Pathway ID | Pathway description | Total pathway size | # Significant genes | FDR-corrected P value |
| --- | --- | --- | --- | --- |
| **Reactome** |  |  |  |  |
| R-HSA-4420097 | VEGFA-VEGFR2 Pathway | 99 | 7 | 0.037 |
| R-HSA-194138 | Signaling by VEGF | 107 | 7 | 0.037 |
| **KEGG** |  |  |  |  |
| KEGG:04611 | Platelet activation | 122 | 8 | 2.5E-03 |
| KEGG:05206 | MicroRNAs in cancer | 158 | 8 | 0.016 |

### Aged COVID-19 patients exhibit increased plasma VEGF protein levels and high peripheral neutrophil to lymphocyte ratio

To further address if our findings were relevant in the human setting of SARS-CoV-2 infection, we stratified COVID-19 patients into aged group (>60 years) and a group of COVID-19 patients <60 years (**Table S5**). We found that with increasing age, there were increased association of disease parameters and comorbidities (**Table S5**). We measured the levels of human plasma proteins levels for IFN-α, IFN-β and IFN-γ. While levels of plasma IFN-α, and IFN-β were below the levels of reliable detection, we found that the COVID-19 patients who were <60 years expressed significantly higher plasma IFN-γ levels when compared to levels in plasma of healthy controls (**Fig. 8A**). Although plasma levels of IFN-γ protein was also increased in aged COVID-19 patient group, levels were not significantly different from healthy controls (**Fig. 8A**). This was in contrast to plasma protein levels of VEGF, which was significantly higher in aged individuals with COVID-19 disease when compared with levels in individuals with COVID-19 disease who were <60 years old (**Fig. 8B**). The increased levels of VEGF in aged COVID-19 patients coincided with significantly increased peripheral neutrophil counts as well as increased peripheral neutrophil to lymphocyte ratios, when compared with both healthy controls and COVID-19 group <60 years old (**Fig. 8C,D**). These results show that plasma protein levels of VEGF and accumulation of peripheral neutrophils is increased in aged individuals with COVID-19 disease, when compared to younger individuals with COVID-19 disease.

**Figure 8.**
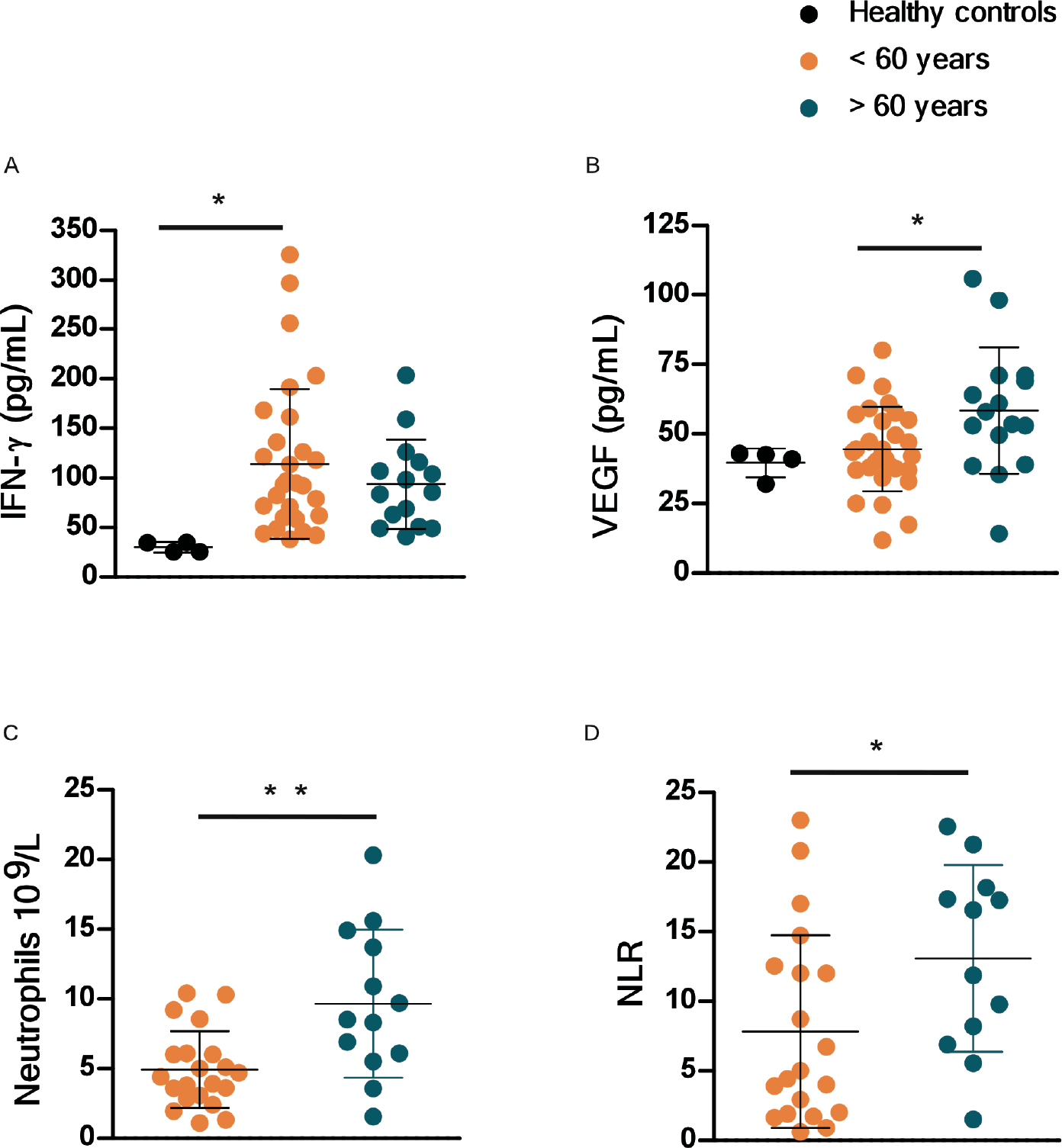
VEGF and peripheral neutrophil counts are higher in old COVID-19 patients. Peripheral blood samples were obtained from a cohort of patients with laboratory-confirmed SARS-CoV-2 infection at hospital admission. Levels of different immune markers were determined by Luminex assay in plasma samples from COVID-19 and healthy volunteer controls. COVID-19 patients were stratified by age as younger than or older than 60 years. (A) Levels of IFN-γ and (B) levels of VEGF proteins were measured in plasma of COVID-19 and healthy controls. Peripheral neutrophil counts (C) and neutrophil to lymphocyte ratio (NLR) values (D) were retrieved from the medical records of COVID-19 patients and compared between age groups.

### Neutrophil degranulation and IFN pathways overlap between COVID-19 and TB disease

Tuberculosis (TB) is a pulmonary granulomatous disease caused by infection with *Mycobacterium tuberculosis*. TB disease in humans and macaques is associated with a neutrophil and IFN signature(*13*). Thus, we next compared and contrasted the transcriptional profile of genes and pathways that are shared by the two diseases, and those that are unique to COVID-19.There was not a substantial overlap between differentially expressed genes in response to COVID-19 and TB. However, of the 97 genes that were commonly upregulated in TB and COVID-19 (**Figure 9A**, **Table S3K**), the Reactome pathway enrichment was well featured in “Neutrophil degranulation”, “Innate immune response”, and “Interferon gamma signaling” (**Figure 9B, Table S4E**). Nearly as many genes (76) had opposite differential expression patterns (upregulated in COVID-19, downregulated in TB), as genes upregulated in both (**Figure 10A**, **Table S3L**). These genes were associated with blood vessel morphogenesis and angiogenesis including leptin receptor (LEPR), TGFβ2 (**Figure 10B, Table S4F**). These results suggest that both TB and COVID-19 share features of neutrophil accumulation of IFN signaling, but that COVID-19 disease immunopathogenesis uniquely features vascularization of the lung.

**Figure 9:**
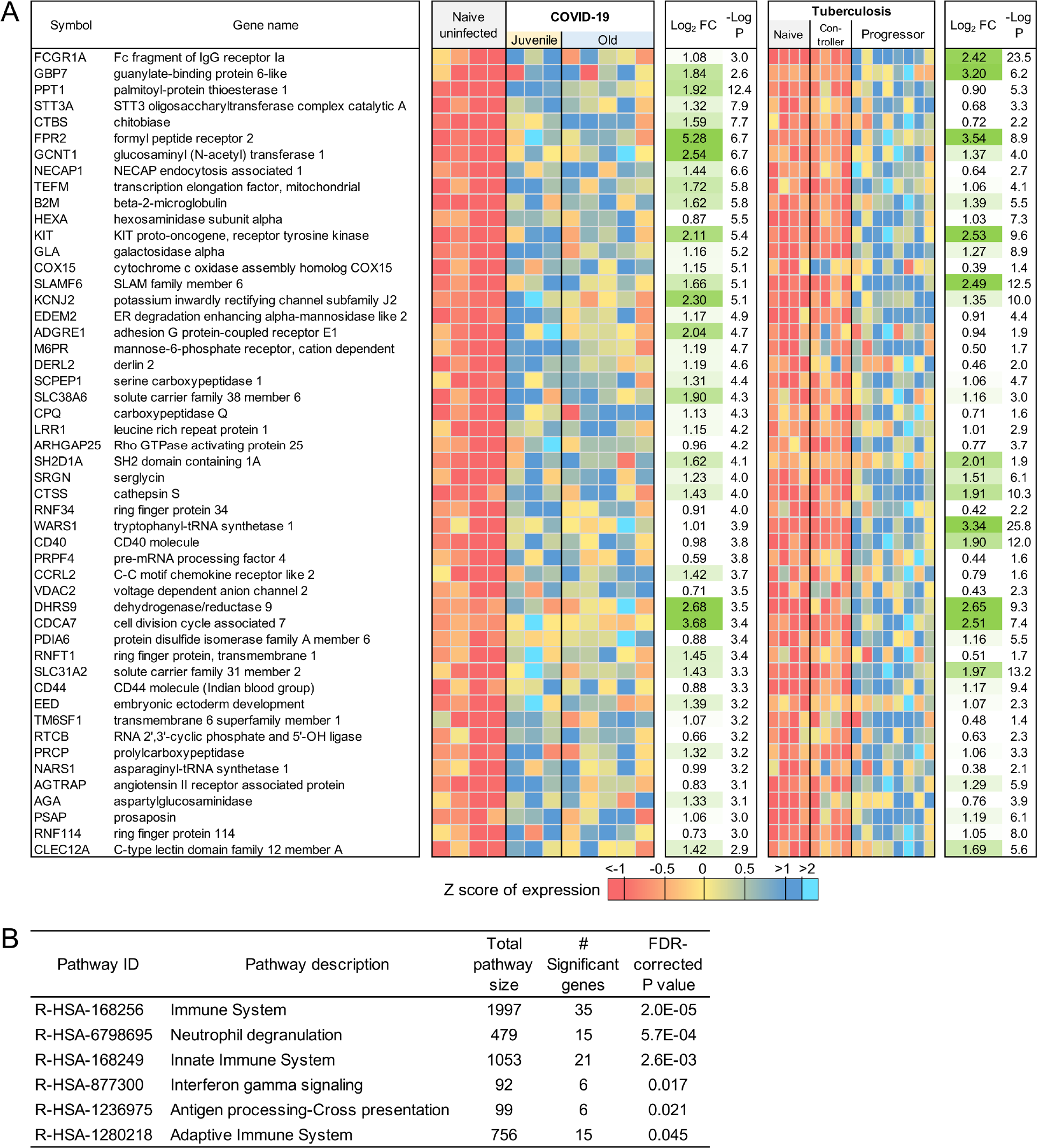
Genes higher in expression during both COVID-19 and TB share common pathways. The top 50 (of 97) most significantly upregulated genes in COVID-19 infected and TB infected macaques. Expression values are visualized by Z scores of normalized expression data (FPKM) per sample, and Log2 Fold Change and −Log P values are from the DESeq2 output. Genes are sorted by P value. (**B**) Significant Reactome pathway enrichment among the 97 genes.

**Figure 10:**
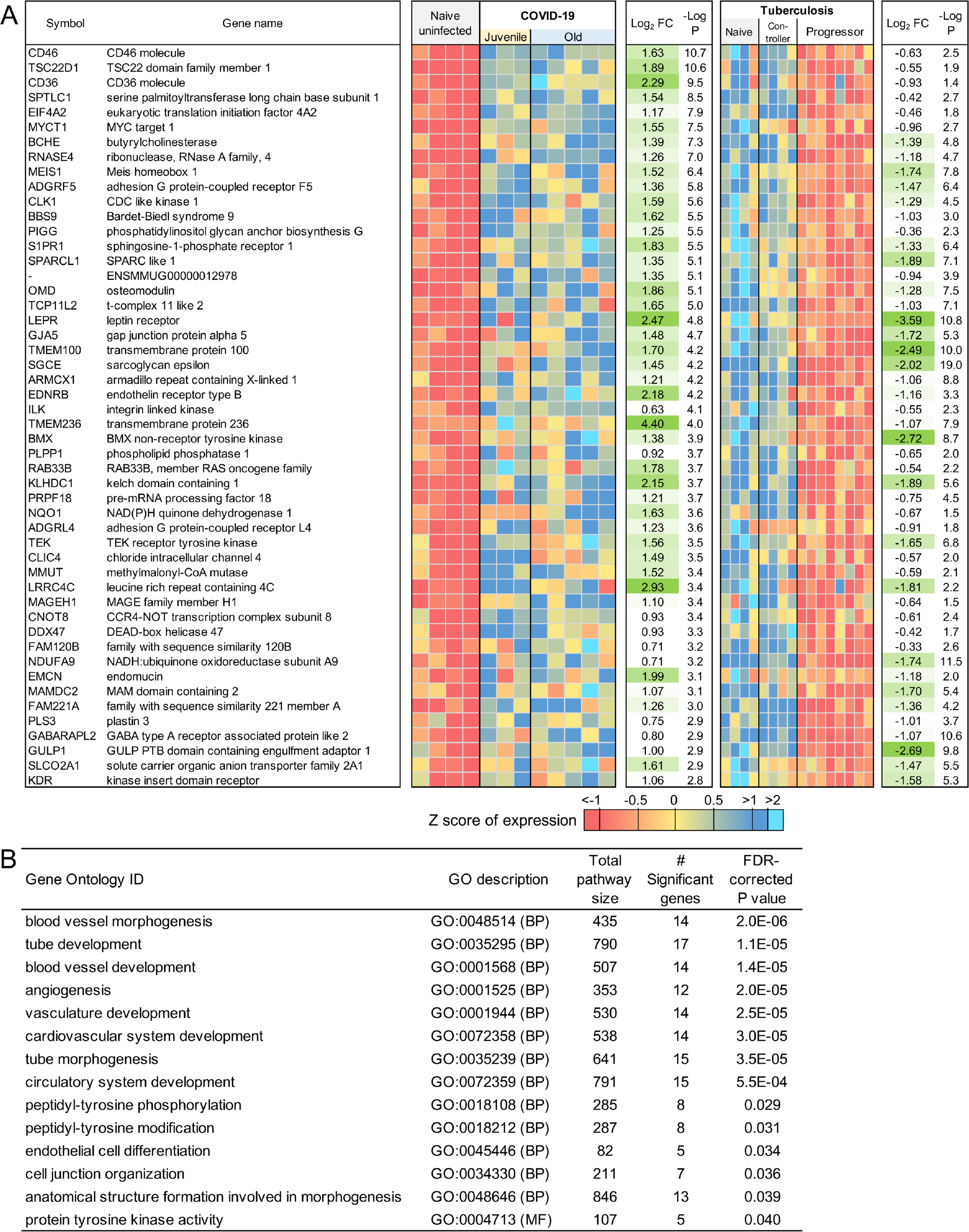
Genes higher in expression during COVID-19 than TB are related to blood morphogenesis pathways. (**A**) The top 50 (of 76) most significantly upregulated genes in COVID-19 infected compared to TB-infected macaques. Expression values are visualized by Z scores of normalized expression data (FPKM) per sample, and Log2 Fold Change and −Log P values are from the DESeq2 output. Genes are sorted by P value. (**B**) Significant Gene Ontology pathway enrichment among the 76 genes.

## DISCUSSION

Lack of understanding of the complexity of COVID-19 immunopathogenesis hampers identification of therapeutic strategies for COVID-19. While studies using immune profiling in COVID-19 patients have shed light on related immune mechanisms of this disease, these have primarily involved peripheral samples obtained from moderate to severe COVID-19 patients, who are generally also older. To overcome these limitations, we have generated a nonhuman primate model (rhesus macaques) of SARS-CoV-2 infection that reflects several features of the immunopathogenesis of human COVID-19, and provides a platform to interrogate the immune pathways that mediate disease versus protection, especially in the context of young versus older hosts. In this study, we show that upregulation of pathways characteristic of neutrophil degranulation and IFN signaling are characteristic of COVID-19 disease in infected hosts. Importantly, the significantly higher induction of genes associated with Type I IFN signaling pathway and Notch signaling in young macaques infected with SARS-CoV-2 is a key determinant that distinguishes them from infected old macaques. Lungs of old macaques infected with COVID-19 however, uniquely feature downregulation of VEGF signaling pathways. Importantly, in PBMCs of humans infected with SARS-CoV-2 we found increased levels of VEGF and peripheral neutrophil counts in individuals >60 years when compared to younger individuals. These results together provide novel insights into the immunopathogenesis of COVID-19 disease, especially from the unique perceptive of age as a contributing factor.

As we learn more about the pathophysiology of COVID-19, it is becoming clear that disease severity is associated with hyperinflammation which in turn induces lung and multiorgan injury and mortality via a cytokine storm (*1, 2, 56*). While therapeutic options that focus on immunomodulatory agents such as corticosteroids are being considered and used, a risk exits that immunomodulators may also inhibit protective pathways. Therefore, a thorough understanding of the host inflammatory responses during SARS-CoV-2 infection is needed before precise immunomodulators can be specifically designed to limit inflammation without regulating protective mechanisms of action. The distinct role of myeloid cells in COVID-19 lung injury and immunopathogenesis is just beginning to be described, and we have clearly shown that neutrophils are intensely recruited to the lung compartment in macaques after SARS-CoV-2 infection (12) (in review). Neutrophils can play a protective role contributing to early antiviral defense (*57*), but also can be pathological due to processes associated with degranulation and lysis, thereby promoting lung inflammation. Consistent with this notion, in current COVID-19 literature, an increased peripheral neutrophil-to-lymphocyte ratio is observed in severe COVID-19 cases, and in some studies is also associated with unfavorable prognosis (*58*). These results in human studies are consistent with our macaque studies that describe neutrophil degranulation as one of the top transcriptional pathways up-regulated in the lungs of COVID-19 macaques when compared to uninfected controls. In this regard, expression of Cathepsin G is northworthy since it is prominent serine protease that amplifies inflammation by stimulating the production of cytokines and chemokines that drive immune cell recruitment to the lung (*59*), and activates metalloproteases to cleave extracellular matrix proteins, thereby promoting neutrophil migration (*60*). Cathepsin G also induces potent chemotactic recruitment of monocytes, neutrophils and antigen presenting cells in addition to promoting endothelial and epithelial permeability (*61*). The latter function of Cathepsin G could be important in enhancing viral invasion to extra-alveolar sites while increased epithelial permeability might also explain the gastrointestinal route of transmission (12) (in review). Additionally, ATP6AP2, causes secretion of inflammatory and fibrotic factors (*41*), CD36, that induces neutrophil apoptosis, and CECAM8 whose cross-linking induces IL-8 production, all of which are highly expressed in COVID-19 diseased lungs. In patients with severe COVID-19, neutrophils express higher frequency of CD66b^+^ neutrophils(*62*). These different genes that are up-regulated as part of the neutrophil degranulation/innate immune response pathways suggest a prominent role for neutrophils that can promote inflammation and a cytokine storm leading to COVID-19 disease pathogenesis. Furthermore, our studies shed light on the importance of the membrane glycoprotein, CD36 in the response to SARS-CoV-2 infection. CD36 is expressed on platelets, macrophages and even epithelial cells. In addition to its well characterized apoptotic function, CD36 is also a receptor for thrombospondin-1 and related proteins and can function as a negative regulator of angiogenesis(*78*). This is particularly important given that angiogenesis is an important feature in patients with COVID-19 and associated ARDS (*79*). CD36 also binds long-chain fatty acids and facilitates their transport into cells, leading to muscle utilization, coupled with fat storage. This contributes to the pathogenesis of metabolic disorders, such as diabetes and obesity and atherothrombotic disease (*79*). A recent single-cell analysis revealed significantly higher CD36 expression in association with ACE2-expressing human lung epithelia cells (*80*). Increased CD36 expression may therefore provide a protective role from extreme lung injury during COVID-19, which is observed in the macaques. Our novel findings that CD36 (as well as other prominent signaling pathways) may be involved in the pathogenesis of COVID-19 has implicaitons for host-direc ted therapy for SARS-CoV-2 infection. In contrast, neutrophils are recruited into the lung very early following macaque infection with SARS-CoV-2(*12*) (in review). Additionally, in lungs of deceased individuals with severe COVID-19 disease neutrophil infiltration occurred in pulmonary capillaries and was accompanied with extravasation of neutrophils into the alveolar space, and neutrophilic mucositis(*63*). In the case of COVID-19, neutrophils could also be a source of excess neutrophil extracellular traps (*64*). Cytokine storm characterized by increased plasma concentrations of IL1β, IL2, IL6, IL7, IL8, IL10, IL17, IFNγ, IFNγ-inducible protein 10, monocyte chemoattractant protein 1 (MCP1), G-CSF, macrophage inflammatory protein 1α, and TNFα seen in severe COVID-19 patients can regulate neutrophil activity by upregulating the expression of chemoattractants that recruit myeloid cells to the lung. These results are also consistent with upregulation of pathways associated with immune and innate signaling, especially IFN signaling. These results together suggest a scenario in the lung where induction of the cytokine storm drives the recruitment of neutrophils, thereby contributing to inflammation. Thus, degranulation of neutrophils and formation of NETs may further promote cytokine responses and inflammation and disease immunopathogenesis.

The IFN response constitutes the major first line of defense against viruses. Recognition of viral infections by innate immune sensors activates both the type I and type III IFN response. While some studies have shown that serum of COVID-19 patients contains increased expression of pro-inflammatory cytokines and chemokines, without detectable levels of type I and III IFNs(*65*), other studies suggest that the IFN response may be delayed. Importantly, elevated IFNs correlate with more severe disease(*66, 67*). However, it is not fully clear if type I IFNs are protective or pathological in COVID-19(*68*). Thus, it is possible that severe infection drives the higher expression of genes in the IFN pathways, but may not lead to viral containment, but instead drives pathological damage. On the other hand, increased induction of type I IFN signaling pathways in SARS-CoV-2 infected macaques, as well as increased induction in juvenile macaques, could support a role for IFN signaling in protection rather than disease progression. Our studies provide data to support the recently proposed hypothesis that that IFN induction may be compromised in older hosts(*68*). When the early IFN response is not optimal to control viral infection, it is possible that delayed or inadequate IFN responses may lead to inflammation mediated damage. Not all animal models, especially mice fully mimic the spectrum of human disease caused by SARS-CoV-2, likely due to the regulatory responses of IFNs on viral entry receptors such as ACE2 which are differentially regulated in humans compared to mice. Further testing the protective versus pathological roles of IFNs in the macaque model with the availability of IFNAR blocking reagents should further clarify the specific role of IFN pathways in COVID-19.

ARDS in influenza, MERS and SARS have been associated with fibrotic irreversible interstitial lung disease(*69, 70*). Pulmonary fibrosis is a recognized sequelae of ARDS(*47*). Pulmonary fibrosis can develop either following chronic inflammation or as a consequence of genetically associated and age-related fibroproliferative process, as in idiopathic pulmonary fibrosis (IPF)(*71*). Fibrosis is the hardening, and/or scarring of tissues due to excess deposition of extracellular matrix components including collagen. Fibrosis is often the terminal result of inflammatory insults induced by infections, autoimmune or allergic responses and others. It is thought that the mechanisms driving fibrogenesis are divergent from those modulating inflammation. The key cellular mediator of fibrosis is the excessive accumulation of fibrous connective tissue (components of the ECM such as collagen and fibronectin) in and around inflamed or damaged tissue. Since a significant proportion of COVID-19 patients develop severe ARDS, it is predicted that a similar outcome of fibrosis will be associated with COVID-19. Also, since the risk factors associated with COVID-19 including increasing age, male and associated co-morbidities coincide with IPF risk factors, it is expected that COVID-19 patients will experience fibrotic lung disease. Despite these associations, there is no evidence currently that “scarring of the lung” experienced by COVID-19 patients is fibrotic or progressive and an outcome of COVID-19 disease post recovery. Therefore, our results provide unique insights into the role of fibrosis during SARS-CoV-2 infection. Most notably, we find significant downregulation of collagen degradation pathways, as well as pathways associated with collagen formation, collagen trimerization and assembly. Furthermore, the role for TGF-β and ECM degradation is well documented in fibrosis. Indeed, the genes associated with these pathways are also significantly down-regulated. These results for the first time provide novel insights into the early pathological events occurring during COVID-19 in the lungs with relevance to underlying immune mechanisms associated with canonical fibrosis pathways. While long term consequences of the pulmonary COVID-19 such as fibrosis remain to be determined, our results on down-regulation of collagen degradation and TGF-β pathways may represent important early events on the lungs of SARS-CoV-2 infected individuals. We speculate that such events may protect individuals from progression to ARDS and fibrosis, while it is possible that in individuals with early activation of collagen degradation progress more severe outcomes may ensue.

Finally, we provide novel insights into the transcriptional regulation of immune pathways that are induced and regulated by age, an important risk factor for COVID-19 disease and outcome. This is a significant component of risk for disease and prognosis of COVID-19. We find higher induction of genes associated with Type I IFN signaling and Notch signaling in the old mecaque. Up-regulation of these significant Type I IFN signaling genes suggest that in a model of self-limited clinical disease in macaques, Type I IFN induction may be differentially regulated by age-associated factors. Age-specific regulation of this pathway has been demonstrated in the murine model of TB(*72*). There is also a well-documented relationship between Notch signaling and viral infections. For example, Human Papilloma Virus and Simian Virus 40 can highjack the cellular machinery, including components of Notch signaling, and these events re associated with cancer progression(*73*). Most studies thus far have only followed SARS-CoV-2 infected macaque for up to two weeks, and it was initially thought that this virus causes acute infection. However, details are now emerging from both animal models(*12*) (in review) and patients, that the virus can persist for longer periods, leading to persistent shedding from tissues, and exhaustion of adaptive responses. While innate and T cell responses are comparable between juvenile and old macaques following infection, SARS-CoV-2 specific antibody is generated at significantly higher levels in the plasma of juveniles, relative to old macaques(*12*) (in review). Since Notch signaling regulates multiple stages of B-cell differentiation and shapes the antibody repertoire(*74*), higher expression of many of the Notch pathway member genes in juvenile macaques may be responsible for the development of stronger antibody responses in these animals, impacting disease progression. Alternatively, it is possible that the differences in Notch signaling and production of virus-specific antibody between jouvenile and old macaques may impact disease progression over a longer period of time, or be particularly relevant in models of co-morbidity, such as diabetes. Similarly, Type I IFN responses are critical for the downstream breadth of antibody production and recognition (*75–77*). Thus, while T cell responses are comparable in juvenile and old macaques, differences in critical signaling pathways uncovered by our RNA-seq analysis potentially explain why juvenile macaques mount significantly stronger antibody responses, and consequently why younger subjects have reduced susceptibility to COVID-19. While this has not been recapitulated in the macaque model, older patients of COVID-19 are more susceptible to progression. This is consistent with increased disease progression when COVID-19 patients were stratified based on age. A previous study found that peripheral VEGF concentrations were significantly higher in COVID-19 patients than in healthy controls(*81*). We also find this effect in our human samples (**Figure 8B**) where people with COVID-19 that are older than 60 years of age have more VEGF protein in their peripheral blood. However, we also find significantly lower levels of VEGF pathway gene transcripts in the lungs of macaques with SARS-CoV-2 infection, especially older macaques (**Figure 6, 7**). Our study further demonstrates that the changes in VEGF signaling may be associated with increasing age rather than just with disease severity. VEGF pathways promote angiogenesis and induce vascular leakiness and permeability. Our results therefore suggest that higher levels of VEGF in the periphery, while a biomarker for COVID-19, may be driven as a compensatory mechanism due to lower levels of VEGF signaling at the site of infection, i.e, the lung. These results further underscore the value of studying responses to SARS-CoV-2 infection in the lung compartment. By uncovering new aspects of the role of these signaling pathway in SARS-CoV-2 infection in the lung compared to the periphery using animal models and human samples, will shed further light on pathways that can be harnessed for therapeutics for COVID-19 disease.

TB and COVID-19 both primarily affect lung function. TB was already one of the leading causes of death due to an infectious disease prior to emergence of COVID-19. In the current scenario the clinical management of both TB and COVID together, particularly in the endemic regions is another rapidly emerging healthcare challenge needing immediate attention. In order to properly address the solution for this emerging crisis a better understanding of the comparative immunological manifestations of both the diseases must be understood. Our results are the first to clearly demarcate the main differences in the manifestation of both the diseases in the alveolar niche. Neutrophil degranulation was one of the most significantly enriched pathways in both the disease conditions and therefore appears as a promising druggable target for efficient management of severe co-morbid TB COVID-19 condition. However, the selective enrichment of angiogenesis and vascular permeability in observed in the lungs of SARS-CoV-2 infected macaques is not seen in models, or patients of TB. These results have the potential to generate additional, specific druggable targets for COVID-19.

Overall, we interrogated transcriptional profiles of lungs from juvenile and old macaques infected with SARS-CoV-2. This study has provided fundamentally new information on the host response in young and old macaques infected with SARS-CoV-2, a model that provides relevant insights necessary for further vaccine and therapeutic development for COVID-19 and a subset of these observations confirmed in human samples with control of SARS-CoV-2 infection as well as COVID-19 disease, and as a function of age.

## Acknowledgements

NHP samples used in this work was derived from studies supported by intramural funds raised by Texas Biomedical Research Institute towards its Coronavirus Working Group, by Regeneron, Inc. (R.C., contract # 2020_004110, in part with federal funds from the Department of Health and Human Services; Office of the Assistant Secretary for Preparedness and Response; Biomedical Advanced Research and Development Authority, under Contract No. HHSO100201700020C). The work described in this manuscript was supported by Washington University in St. Louis (S.A.K) for COVID-19 research, as well as and NIH award # R01AI123780 to S.A.K, M.M. and D.K., R01AI134236 to S.A.K. and D.K. and a COVID-19 supplement to it., and by institutional NIH awards P51OD111033 and U42OD010442 to the SNPRC, Texas Biomedical Research Institute. J.A.P-C was supported by the National Council of Science and Technology of Mexico to achieve (CONACYT) his PhD degree (CONACyT-CVU 737347). The current study was supported by institutional research funds of INER and by research contracts: SECTEI/050/2020, Secretaría de Ciencia, Tecnología e Innovación de la Ciudad de México (SECTEI CDMX); FORDECYT/10SE/2020/05/14-06 and FORDECYT/10SE/2020/05/14-07 from the Fondo Institucional de Fomento Regional para el Desarrollo Científico y Tecnológico y de Innovación (FORDECYT), Consejo Nacional de Ciencia y Tecnología (CONACYT). These funders had no role, however, in the design and execution of the experiments and the interpretation of data. The views expressed here are those of the authors and do not necessarily represent the views or official position of the funding agencies. The authors declare that no other financial conflict of interest exist.

## Author Contributions

B.A.R., M.A., D.S., J.C., B.S., J.M., O. G, J.A.C-P., L.A.J-A., T.S.R-R., J.Z. carried out experiments, analysed data; J.Z., L.S.S., J.T., R.C., M.M., D.K., and S.A.K designed the study, provided funding or reagents; M.A., B.A.R., D.K., and S.A.K wrote the paper; all authors read, edited and approved the manuscript.

## Supplementary Tables

**Table S1:** Read processing and mapping statistics, and download accessions for all RNA-seq samples.

**Table S2**: Fragment counts, relative gene expression levels, gene annotations, and differential expression data for every macaque gene.

**Table S3:** Complete lists of significantly differentially expressed gene sets of interest (including gene names, relative expression data, fold change and P values). Gene sets include: (**A**) 1,026 genes significantly up-regulated with COVID-19 vs Naive, (**B**) 65 “neutrophil degranulation” (R-HSA-6798695) genes significantly up-regulated during COVID-19, (**C**) 162 “neutrophil degranulation” (R-HSA-6798695) genes significantly up-regulated during COVID-19, (**D**) 1,109 genes significantly down-regulated with COVID-19 vs Naive, (**E**) 14 “collagen degradation” (R-HSA-1442490) genes significantly up-regulated during COVID-19, (**F**) 14 “Signaling by TGF-beta Receptor Complex” (R-HSA-170834) genes significantly up-regulated during COVID-19, (**G**) 86 genes significantly up-regulated with COVID-19 vs Naive only in Juvenile macaques, (**H**) 96 genes significantly down-regulated with COVID-19 vs Naive only in Juvenile macaques, (**I**) 97 genes significantly up-regulated with COVID-19 vs Naive only in Old macaques, (**J**) 160 genes significantly down-regulated with COVID-19 vs Naive only in Old macaques, (**K**) 97 genes significantly up-regulated by both COVID-19 and TB and (**L**) 76 genes significantly up-regulated by COVID-19 but down-regulated by TB.

**Table S4**: Significant functional enrichment for Reactome, KEGG and Gene Ontology pathways, among differentially gene sets of interest. Gene sets include: (**A**) 1,026 genes up-regulated in COVID-19 vs Naive, (**B**) 1,109 genes down-regulated in COVID-19 vs Naive, (**C**) 86 genes signficantly up-regulated by COVID-19 only in Juvenile macaques, (**D**) 160 genes signficantly down-regulated by COVID-19 only in Old macaques, (**E**) 97 genes significantly up-regulated by both COVID-19 and TB, and (**F**) 76 genes significantly up-regulated by COVID-19 but down-regulated by TB.

**Table S5**. Clinical characteristics and lab parameters of COVID-19 patients. Clinical and demographic data were retrieved from the medical records of all participants. These data included age, gender, anthropometrics, comorbidities, symptoms, triage vital signs, and initial laboratory test results.

